# Coordination of CcpA and CodY regulators in Staphylococcus aureus USA300 strains

**DOI:** 10.1101/2022.05.25.493525

**Authors:** Saugat Poudel, Ying Hefner, Richard Szubin, Anand Sastry, Ye Gao, Victor Nizet, Bernhard O. Palsson

## Abstract

The complex crosstalk between metabolism and gene regulatory networks makes it difficult to untangle individual constituents and study their precise roles and interactions. To address this issue, we modularized the transcriptional regulatory network (TRN) of the Staphylococcus *aureus* USA300 strain by applying Independent Component Analysis (ICA) to 385 RNA sequencing samples. We then combined the modular TRN model with a metabolic model to study the regulation of carbon and amino acid metabolism. Our analysis showed that regulation of central carbon metabolism by CcpA and amino acid biosynthesis by CodY are closely coordinated. In general, *S. aureus* increases the expression of CodY-regulated genes in the presence of preferred carbons sources such as glucose. This transcriptional coordination was corroborated by metabolic model simulations that also showed increased amino acid biosynthesis in the presence of glucose. Further, we found that CodY and CcpA cooperatively regulate the expression of ribosome hibernation promoting factor, thus linking metabolic cues with translation. In line with this hypothesis, expression of CodY-regulated genes is tightly correlated with expression of genes encoding ribosomal proteins. Together, we propose a coarse-grained model where expression of *S. aureus* genes encoding enzymes that control carbon flux and nitrogen flux through the system is coregulated with expression of translation machinery to modularly control protein synthesis. While this work focuses on three key regulators, the full TRN model we present contains 76 total independently modulated sets of genes, each with the potential to uncover other complex regulatory structures and interactions.

**Importance:** *Staphylococcus aureus* is a versatile pathogen with an expanding antibiotic resistance profile. The biology underlying its clinical success emerges from an interplay of many systems such as metabolism and gene regulatory networks. This work brings together models for these two systems to establish fundamental principles governing the regulation of S. aureus central metabolism and protein synthesis. Studies of these fundamental biological principles are often confined to model organisms such as *Escherichia coli*. However, expanding these models to pathogens can provide a framework from which complex and clinically important phenotypes such as virulence and antibiotic resistance can be better understood. Additionally, the expanded gene regulatory network model presented herein can deconvolute the biology underlying other important phenotypes in this pathogen.

## Introduction

Metabolism plays an integral role in infection and antimicrobial resistance (AMR) in the leading human bacterial pathogen *Staphylococcus aureus*. Metabolic requirements specific to infection, intracellular persistence, biofilm formation, and colonization are rapidly being uncovered^1–6^. Furthermore, the central role of metabolism in AMR and persistence is also coming into view, adding to the complexity of known AMR mechanisms^7–9^. The complex metabolic circuits and responses underlying these phenomena are nevertheless difficult to unravel. Even relatively well-understood systems such as *S. aureus* central carbon metabolism can be difficult to fully map, as they are layered with multiple levels of gene regulation, post-translational and biochemical controls, and unexpected molecular interactions ^1,10–12^. Some of these complexities can be captured by genome-scale metabolic models (GEMs) that allow rapid query of metabolic complexities through simulations of metabolic flux states, knock-out experiments, multi-strain metabolic comparisons, and calculation of metabolic characteristics^13,14^. Alternatively, coarse-grained modeling of metabolism attempts to peer beyond the detailed complexity and discover the general principles governing the biological systems of interest^15–17^. In the present work, we took guidance from a coarse-grained model proposed in *Escherichia coli* coupled with genome scale analyses of *S. aureus* transcriptional regulation and metabolism to uncover a similar staphylococcal system that balances resource allocation between carbon and nitrogen metabolism^15,17^.

Biological trade-offs represent an optimization frontier, where the cell must strike a balance between its multiple objectives and their limitations^15,18^. Signatures of these balancing acts can be found in transcriptomes and become apparent when their architecture is viewed at systems level^19^. We previously described one such trade-off and its transcriptional imprint using independent data sets from Gram-negative *E. coli* and Gram-positive *S. aureus*—in which a balance was observed between genes regulated by stress-associated sigma factors and growth-associated translation machinery^20,21^. Here, we expand significantly beyond those observations to describe a trade-off between carbon and nitrogen metabolism in strains of the globally disseminated, hypervirulent *S. aureus* USA300 lineage.

We first greatly expanded on our previous model of transcriptional regulation in USA300 strains to incorporate all publicly available RNA sequencing data from the Sequence Reads Archive (SRA)^21^. Models were then generated by applying independent component analysis (ICA), which calculates independently modulated sets of genes (iModulons) and their activities present in the input RNA sequencing samples. iModulons represent sources of signals in the expression data, with transcriptional regulators being the most common source. Our model showed that the activities of two global metabolic regulators, CcpA and CodY, which play critical roles in central carbon and nitrogen metabolism respectively, are negatively correlated against one another. This negative correlation pointed to a condition-specific reallocation of resources towards different metabolic subsystems. GEMs fitted with metabolomics data confirmed the inferences made from the transcriptomic data. Furthermore, GEMs revealed specific metabolic interfaces where coordination of metabolism by the two regulators is required for optimal biomass production, including glutamate dehydrogenase and the folate cycle. Placing genes from CodY and CcpA-associated iModulons onto the metabolic map demonstrated that they did not share any metabolic reactions, but coregulated expression of a gene encoding ribosome hibernation factor. In light of these observations, we propose a model whereby CcpA and CodY coordinate gene expression for carbon metabolism, nitrogen metabolism and translation, thus modularly controlling protein production at specific stages.

## Results

### Expanding the USA300 iModulons using RNA-sequencing data from SRA database

Our previous work outlined 29 iModulons for USA300 strains that were generated from 108 in-house RNA-sequencing data^21^. To expand the previous iModulons coverage of the TRN, we took advantage of the rapidly growing, publicly-available *S. aureus* RNA sequencing samples (**Figure S1**). We queried the Sequence Reads Archive (SRA) for all available USA300-specific RNA sequencing data and combined it with 64 newly generated samples. Of the 576 sequencing samples available, 385 passed the stringent QC/QA pipeline and were therefore incorporated into the new model (see Methods). The final set of samples contained data from at least 7 different USA300 isolates, 4 growth phases (exponential, stationary, biofilm and infection) and 10 base media (**Figure S2**).

Before applying ICA, we normalized the log-transformed Transcripts per Million (log-TPM) data to a project specific control condition. This reduced batch-specific variation in the data and reduced the presence of iModulons not associated with biological signals. Principal component analysis of the log-TPM data showed that normalized samples tended to cluster with media types and growth phases rather than by data source (**Figure 1a**). For example, data from *S. aureus* grown to late-log phase in SCFM2 (Synthetic Cystic Fibrosis Sputum Medium 2) and to stationary phase in Chemically Defined Medium (CDM) did not cluster together, despite being from the same bioproject.

**Figure 1.**
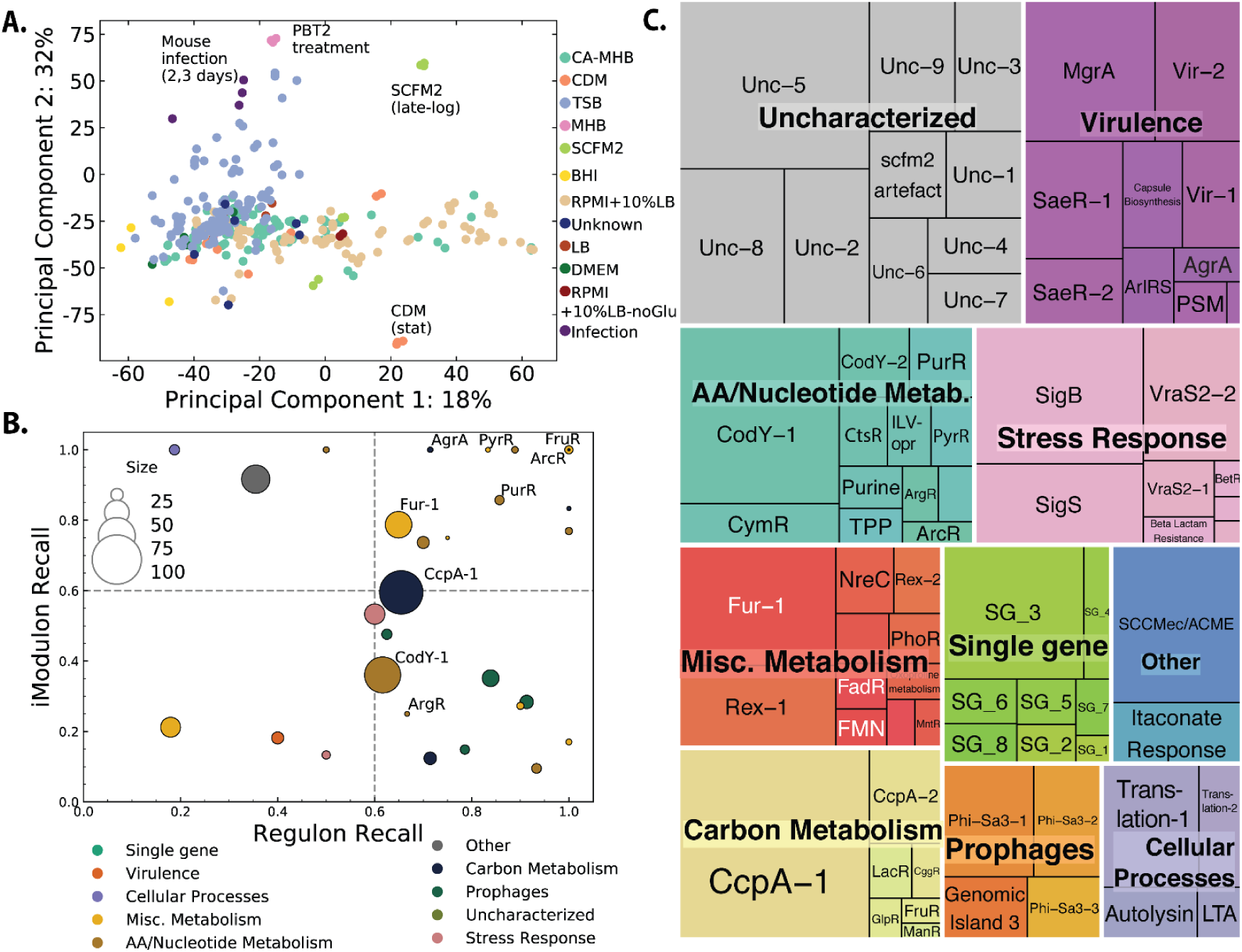
The updated iModulons for USA300 strains. A) 385 RNA-sequencing samples from diverse growth conditions were used to generate the expanded USA300 iModulons. The samples were normalized to project specific control conditions to reduce signal from batch effect. B) iModulons were labeled based on significant association with other published regulons. C) Treemap of iModulon names, size (gene content) and types in the current model after manual curation. The size of the boxes correspond to the number of genes in the iModulon.

Application of ICA to this normalized expression data resulted in 76 independent components, and genes with high absolute weightings within each component were assigned to a corresponding iModulon. These enriched iModulon genes were then compared with existing literature of predicted regulons in *S. aureus*. Those iModulons that had significant overlap with other predicted regulons were named after the associated regulator (**Figure 1b**). Lastly, some iModulons with no known regulators, but associated other biological processes (e.g. prophages, translation) were manually curated. In total, we labeled 60 of the 76 iModulons with either a regulator or a biological process (**Figure 1c**). The remaining uncharacterized iModulons represent signals in the *S. aureus* transcriptome with currently unknown origins, thus providing a road map to discovery of missing parts of the known TRN. In addition to the structure of each iModulon, the activities of each of the 76 iModulons in the 385 input samples were also calculated. The activity represents the role each iModulon (and the associated regulator, if known) in shaping the transcriptome in the given sample. Higher iModulon activity represents higher expression level of genes with positive weightings in the iModulon and lower expression of genes with negatively weighted genes^20^.

### CcpA and CodY iModulon activities highlight balance of carbon and nitrogen metabolism

Cumulatively, the 70 iModulons captured ∼70% of the variance in the input transcriptomic data. The CodY-1, CcpA-1 (henceforth referred to as simply CodY and CcpA iModulons, respectively) and Translation iModulons explained the most variation in the data (**Figure 2a**). CcpA is the catabolite repressor protein in firmicutes that represses genes involved in alternate carbon utilization as well as other central carbon metabolic pathways such as the Tricarboxylic acid (TCA) cycle in the presence of high concentrations of glucose. CodY, on the other hand, globally represses the genes required for amino acid biosynthesis in response to high branched chain amino acid (BCAA) or GTP concentrations. Lastly, the Translation iModulon almost entirely consists of ribosomal genes (e.g. *rplK, rplA etc*.) and genes involved in translation such as *infA* and *fusA* which encode translation initiation factor IF-1 and elongation factor G respectively(**Figure 2b**). This iModulon has been enriched in almost all bacteria and archaea for which iModulons have been calculated ^20,22–25^.

**Figure 2.**
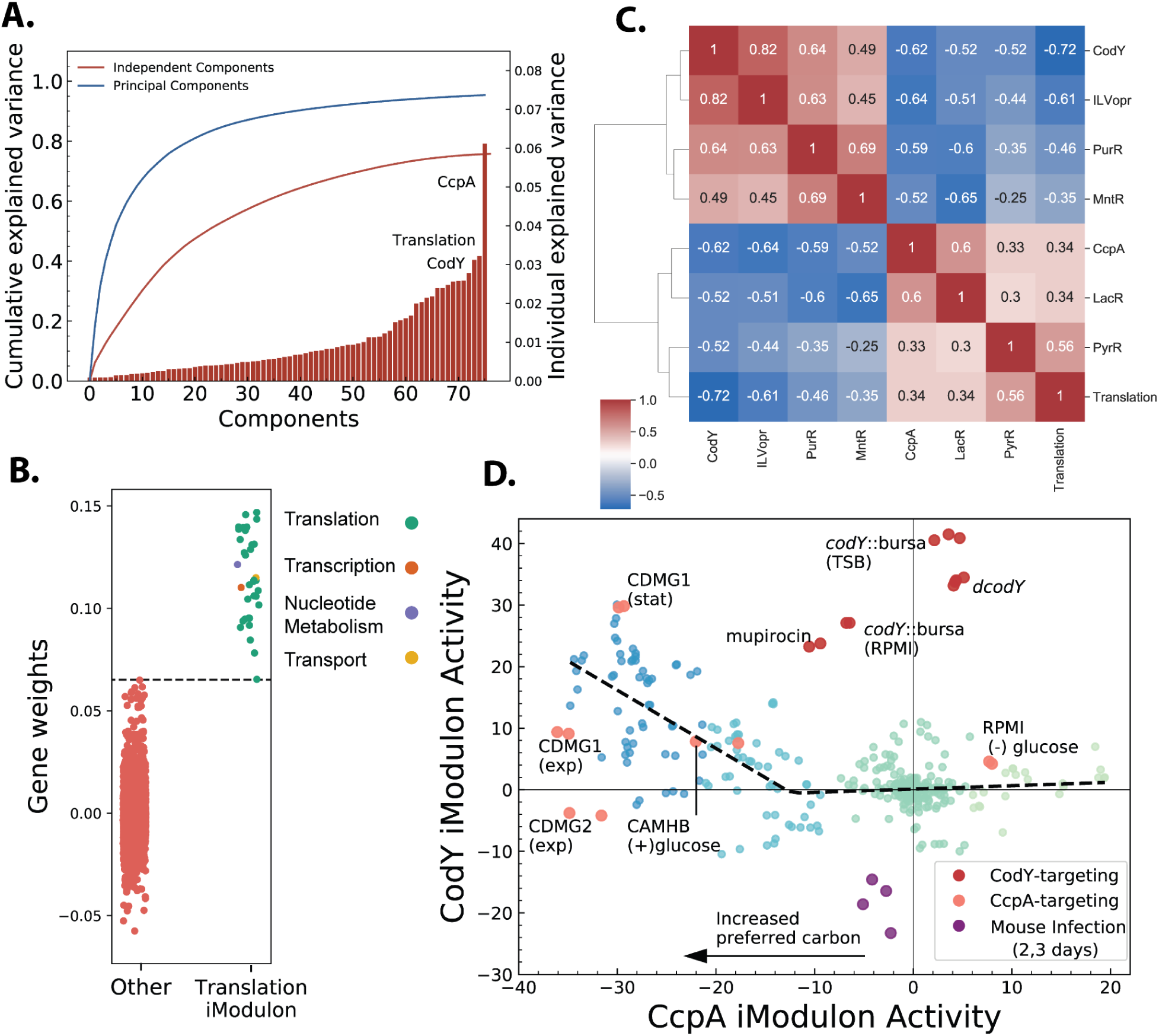
Coordination of metabolic iModulons in USA300 strains. A) Explained variance of each of the iModulons; CcpA, Translation and CodY iModulons explain the most variance in the transcriptome data. B) Translation iModulon gene weightings shows almost all genes enriched in the iModulon are associated with translation. C) Translation iModulon gene weightings shows almost all genes enriched in the iModulon are associated with translation. D) Activity of CcpA and CodY iModulons across all USA300 samples. Inactivation of CodY does not alter CcpA activity but decrease in CcpA activity leads to increase in CodY activity. This asymmetric relationship suggests that CcpA works upstream of CodY. Abbreviations: stat-stationary; exp-exponential; RPMI-Roswell Park Memorial Institute medium; TSB- Tryptic Soy Broth.

Interestingly, activities of these three iModulons were highly correlated across all samples (**Figure 2c**) and formed a large cluster along with other metabolic iModulons (**Figure S3**). Along with CodY, CcpA, and Translation iModulons, activities of ILVopr (iModulon containing the operon with isoleucine, leucine, and valine biosynthesis genes), MntR, LacR and PurR iModulons were also highly correlated. Correlation of CcpA with LacR simply reflects the catabolite repression of lactose utilization genes by the regulator CcpA. Similarly, ILV operon is regulated globally by CodY and locally by leucine attenuator ^26^. This multi-layer regulation likely explains why this operon formed its own iModulon whose activity was closely correlated with CodY. The MntR iModulon contains genes required for manganese uptake, and its coordinated activity with CcpA confirms the association of manganese concentration with glycolytic flux^27^.

The correlated activity of CcpA and CodY iModulons suggested that *S. aureus* carefully coordinates its central carbon and nitrogen metabolism (**Figure 2d**). Close examination of the activities of these two iModulons showed a biphasic relationship. In conditions with preferred carbon sources, and therefore low CcpA iModulon activity, CodY activity generally increased. This effect was observed when glucose was added to both a complex medium (Cation-Adjusted Mueller Hinton Broth, or CA-MHB) and to a defined medium (Chemically Defined Medium or CDM1). Other conditions without explicitly controlled glucose levels that showed low CcpA activity still had concomitant high CodY activity, suggesting that this effect was not glucose specific. In conditions with already low CodY activity however, removal of glucose (RPMI (-) glucose; substituted with maltose) did not lead to further change in CodY activity, creating the second phase of the trade-off plane.

On the other hand, increase in CodY iModulon activity did not necessarily lead to decrease in CcpA activity (**Figure 2d**; red markers). Samples from *codY* interrupted strains in several different projects showed minimal effect on CcpA iModulon activity. These samples fell well outside of the CcpA-CodY trade-off line (**Figure 2d**; gray dashed lines). Similar effects can also be observed in samples treated with sub-inhibitory concentration of mupirocin. Mupirocin activates the stringent response in *S. aureus* which leads to conversion of GTP to ppgpp and subsequent derepression of CodY regulon^28^. As change in CcpA activity leads to change in CodY activity but not necessarily vice-versa, this data suggests that CcpA works ‘upstream’ of CodY.

### Metabolic modeling confirms coupling of CcpA and CodY activities

To independently confirm the metabolic interaction between CodY and CcpA, we used a previously published USA300 strain specific genome-scale metabolic model (GEM) ^29^. GEMs are curated mathematical models of an organism’s metabolic network that can be used to simulate, study, and design the metabolic pathways using a wide range of Constraints Based Reconstruction and Analysis (COBRA) tools^14,30^.

One such method, parsimonious Flux Balance Analysis (pFBA), can be used to calculate metabolic flux state that optimizes a phenotype while minimizing total metabolic flux in a given condition^13,31^. pFBA thus represents a parsimonious use of the metabolic proteome. Here, we used pFBA to determine the metabolic flux states that maximize *S. aureus* biomass production given the measured uptake and secretion rates of various amino acids and sugars in Chemically Defined Medium (CDM) and CDM + glucose (CDMG)^10^. In agreement with increased CodY iModulon activity in CDMG, total flux through reactions catalyzed by enzymes that are encoded in CodY iModulon genes (“CodY reactions” for short), increased from 3.9 mmol/gDW/hr to 5.5 mmol/gDW/hr in presence of glucose (**Figure 3a**).

**Figure 3.**
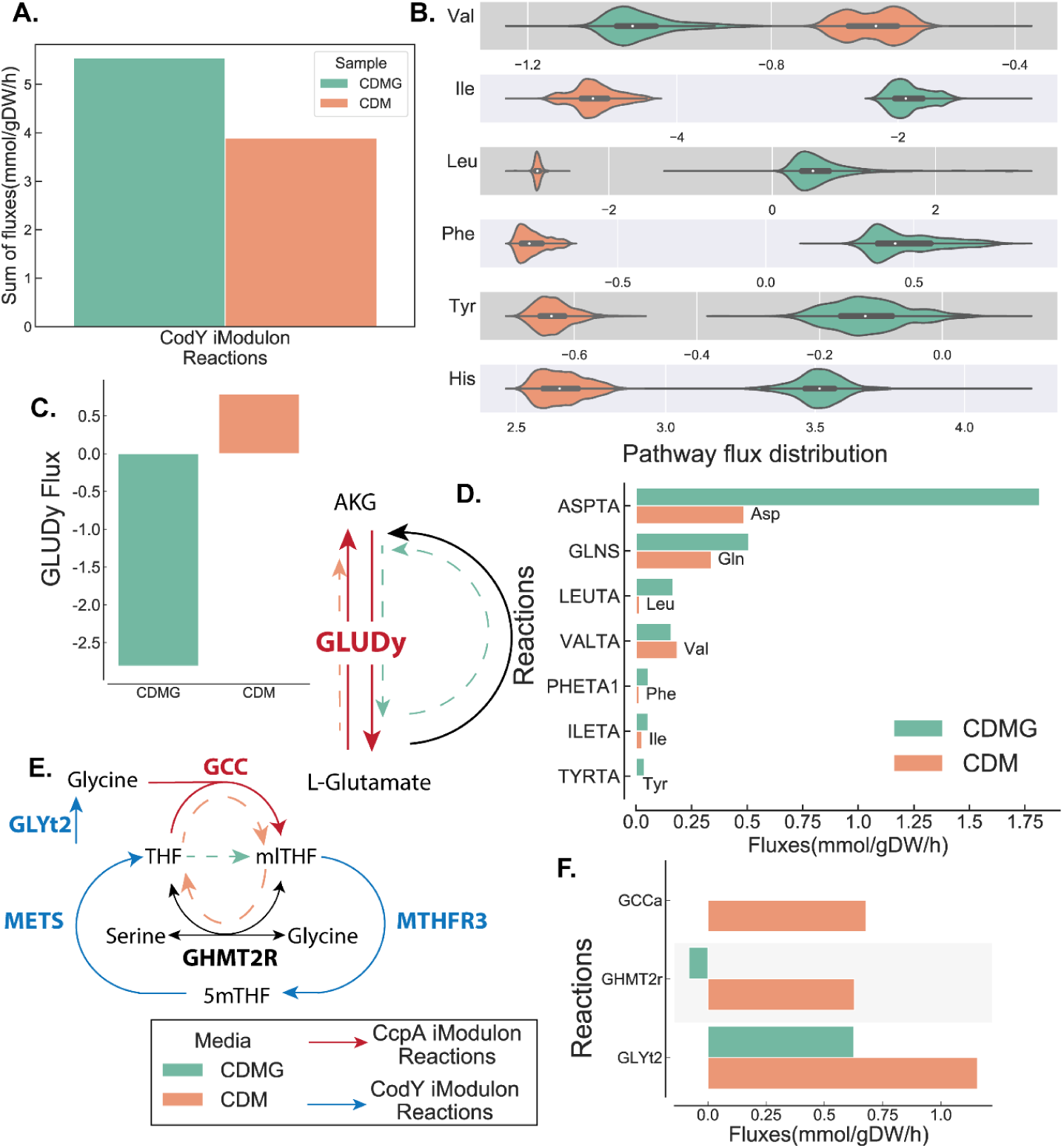
Metabolic modeling shows coordination of CcpA and CodY fluxes. A) Sum of sampled fluxes through CodY and CcpA reaction shows increased flux through CodY in CDMG. B) Sampled fluxes through several amino acid biosynthesis pathways also show increased flux in CDMG. Definition of the biosynthetic pathways is found in Methods. C) Flux through GLUDy reaction changes direction when glucose is added. D) Amino acids generated by accepting amine groups from L-glutamate. L-glutamate is converted to akg in the process and regenerated by GLUDy. E) Metabolic map of folate cycle where CcpA and CodY regulated metabolism intersect. F) Flux through reactions in folate cycle in CDM and CDMG.

pFBA however, gives an exact optimal solution and therefore does not account for variations or errors in input uptake data. We addressed this issue by sampling the CDM and CDMG specific models, which gives a distribution of feasible fluxes in each of the respective conditions. We then mapped the condition-specific flux distributions to various amino acid biosynthetic pathways. For simple interpretation, we excluded amino acids that serve as intermediates for biosynthesis of other amino acids (e.g. glutamine, glutamate and serine) and included only those amino acid for which unique biosynthetic pathways could be defined (see Materials and Methods). Confirming pFBA analysis, 5 out of the 6 amino acid biosynthetic pathways had increased flux in CDMG when compared to CDM (**Figure 3b**). The results of these two TRN-agnostic metabolic modeling methods are in agreement with our observation that CodY iModulon activity increases in the presence of glucose.

### Transcriptional coordination of CcpA and CodY are likely due to flux coupling at metabolic interfaces

CcpA and CodY iModulons contained 110 and 86 genes respectively. Most of these genes are involved in central carbon and amino acid metabolism. Despite the large iModulon sizes and close metabolic proximity of the regulated genes, the two iModulons did not share any genes encoding metabolic enzymes. The correlation in iModulon activities however, suggested that CcpA reactions and CodY reactions must be coordinated at a metabolic level. Using the USA300 GEM, we looked for this coordination at the metabolite interface between CcpA and CodY reactions, i.e., metabolites that are involved in both CcpA and CodY reactions.

We found these metabolic interfaces by systematically identifying all metabolites in USA300 GEM that can be found in both CodY and CcpA reactions. After taking out ‘non-specific’ metabolites and cofactors (e.g., ATP, H _2_O, NADH, etc.), we were left with 22 metabolites at the interface (**Supplementary Table 1**). While some of these metabolites like pyruvate, glutamate and oxaloacetate are expected, as they play a crucial role in both carbon and nitrogen metabolism, other metabolites like N-Succinyl-2-L-amino-6-oxoheptanedioate and tetrahydrofolate (THF) are less understood in the context of this trade-off. To further understand how change in simulated flux through CcpA and CodY reactions in CDM and CDMG altered these key metabolic interfaces, we mapped the pFBA solution fluxes from each media to the reactions around two of these interfaces-glutamate and methylTHF.

The glutamate-alpha ketoglutarate (αkg) link is a closely studied interface in *S. aureus* that connects amino acid and central carbon metabolism ^5,10^. The main enzyme at the interface, glutamate dehydrogenase (GLUDy) reversibly iterconverts αkg and glutamate and is encoded by *gudB* gene, a constituent of the CcpA iModulon. However, this interconversion also acts as an amine group donor or acceptor to 3 CcpA reactions and 8 CodY reactions (**Supplementary Table 1**). In glucose free CDM, pFBA solution was consistent with previous observation showing proline is converted to αkg via glutamate and eventually fuels gluconeogenesis^10^. However, in CDMG, the flux through GLUDy changes direction and catalyzes conversion of αkg to glutamate instead (**Figure 3C**). This makes up ∼98% of total flux that consumes αkg. The glutamate in turn acts as an amine group donor for biosynthesis of various amino acids and accounts for ∼80% of total flux generating αkg in CDMG (**Figure 3D**). pFBA solution of this interface therefore shows that in absence of glucose, GLUDy reaction converts glutamate to αkg to fuel gluconeogenesis but in the presence of glucose it converts αkg to glutamate to fuel amino acid biosynthesis.

The folate cycle represents another metabolic interface of CcpA and CodY reactions. The folate cycle is required for one carbon metabolism, nucleotide biosynthesis and amino acid metabolism and the pathway leading up to the cycle is the target of sulfonamide class antibiotics^*32*^. The folate cycle consisted of 2 CodY reactions - MTHFR3 and METS (methionine synthase) - and one CcpA reaction - GCCabc (glycine cleavage complex) (**Figure 3E**). In CDM, tetrahydrofolate (THF) is converted to 5,10-methylenetetrahydrofolate (mlTHF) by GCCabc reaction which cleaves glycine in the process (**Figure 3F**). THF is then regenerated from mlTHF by GHMT2r reaction which also consumes glycine and generates serine. This consumption of glycine in folate cycle by CcpA reaction is coupled with increased transport of glycine by CodY regulated GLYt2. However, in CDMG where CcpA iModulon activity is low, there is no flux through the CcpA reaction, GCCabc. Instead, GHMT2r runs in ‘reverse’ to convert THF from mlTHF consuming serine and generating glycine instead. Together, combining iModulon structure with metabolic simulation demonstrates how *S. aureus* coordinates flux through CcpA and CodY iModulon reactions at these key metabolic interfaces, despite not sharing any genes at regulatory level.

### The expression of translation associated genes are responsive to CcpA and CodY activities

While CcpA and CodY iModulons do not share any metabolic genes, *hpf*, which encodes ribosomal hibernation promoting factors (HPF), is a gene found in both iModulons. HPF is a small peptide that dimerizes 70S ribosomal subunits to form inactive 100S subunits^33,34^. It plays an important role in stress response, nutrition limitation and protects ribosomal pools from degradation^35–37^. Previous studies in *S. aureus* have shown that SigB and CodY regulate *hpf* expression in response to heat and nutritional stress^35^. iModulon structure confirms the role of the CodY and suggests and additional layer of control by CcpA.

ChIP-exo data from our previous work found two CodY binding sites in the regulatory region of the *hpf* gene (**Figure 4b**)^38^. To confirm the role of CcpA in *hpf* expression, we searched for the catabolite repressor element motif (WTGNNARCGNWWWCAW) from *Bacillus subtilis* in the same region^39^. A matching motif was found in the region between the two CodY binding peaks (q-val=0.00905). This architecture, with two CodY binding sites flanking the CcpA binding site, is also found in the regulatory region of *B. subtilis* BCAA operon where both regulators contribute to the expression of the operon genes^40^. The signal from expression data and the presence of binding motifs suggests that CcpA regulates *hpf* along with previously identified regulators CodY and SigB.

**Figure 4.**
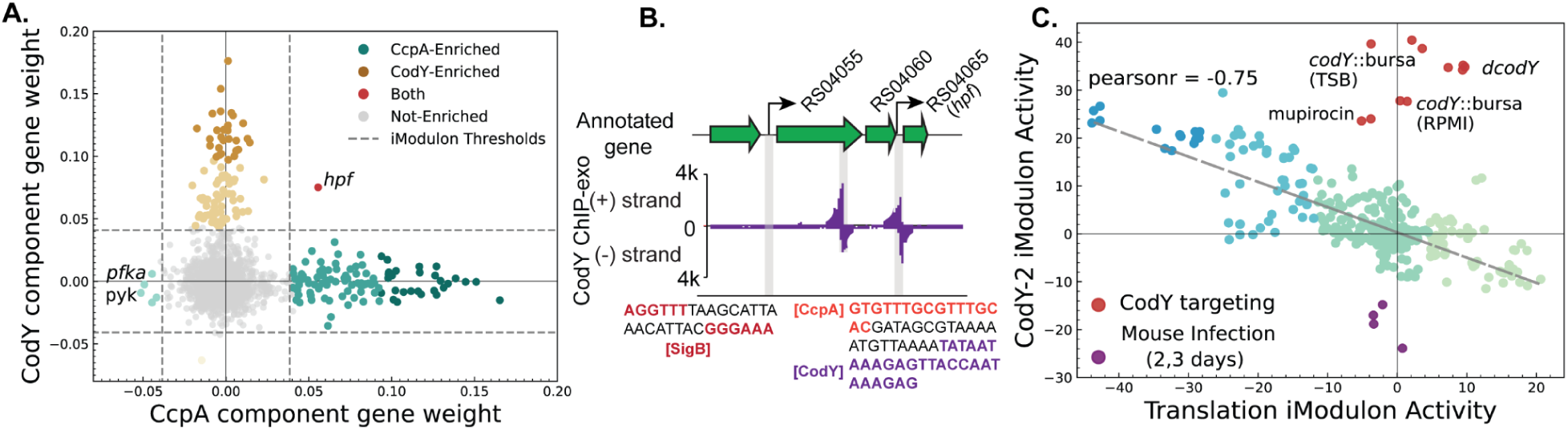
Coordination of translation with metabolism. A) Gene weights in CcpA and CodY iModulons shows that only the hpf gene is a part of both iModulons. B) Upstream region of hpf gene with its two alternative transcription start sites. Two CodY binding sites were detected by ChIP-exo (purple bars). The previously recognized SigB(red) and Cody(purple) binding sites and newly proposed CcpA (orange) binding site are highlighted. C) The negative correlation between CodY and Translation iModulon suggests coordination of metabolism and translation in S. aureus.

In addition to coordinated regulation of translation associated *hpf* gene, CodY activity was also strongly correlated with Translation iModulon activity. In contrast, CcpA and Translation iModulon activities showed little correlation between them (**Figure S4**). Similar to CcpA and CodY activity correlation, *codY* knockout and stringent response activation by mupirocin also disrupted correlation with Translation iModulon (**Figure 4c**). This also suggested that the signal controlling Translation iModulon gene expression also works ‘upstream’ of CodY as interruption of CodY had little effect on Translation iModulon activity. While the coordination of the two iModulon activity is apparent, we were unable to further interrogate the nature of this relationship since the signal behind the Translation iModulon is yet to be identified.

## Discussion

Based on the data presented here, we propose a coarse-grained model of transcriptional regulation of metabolism involved in protein synthesis in *S. aureus* USA300 strains (**Figure 5**). It is motivated by the model of proteome coordination in *E. coli* and extends its principles to non-model pathogenic organism^15^. The coarse-grain model simplifies metabolism underlying protein synthesis into three steps; **(1)** the generation of precursors from carbon source, **(2)** biosynthesis of amino acids from precursors or direct transport from the medium and **(3)** synthesis of peptides from amino acids via translation. The generation of precursors from carbon sources is largely regulated by CcpA (purple arrow). CcpA represses alternate carbon sources (including amino acids such as proline, glutamine and aspartate) in the presence of preferred carbon (such as glucose) and regulates other key aspects of central metabolism such as gluconeogenesis and TCA cycle that are necessary to generate various precursors^1,10,42–44^. The precursors in our model are represented by the metabolites at the CcpA-CodY interface derived from the USA300 GEM (**Supplementary Table 1**).These precursors are then converted to amino acids via CodY regulated gene products (green arrow) and polymerized by ribosomes into proteins (light blue arrow)^38,44,45^.

**Figure 5.**
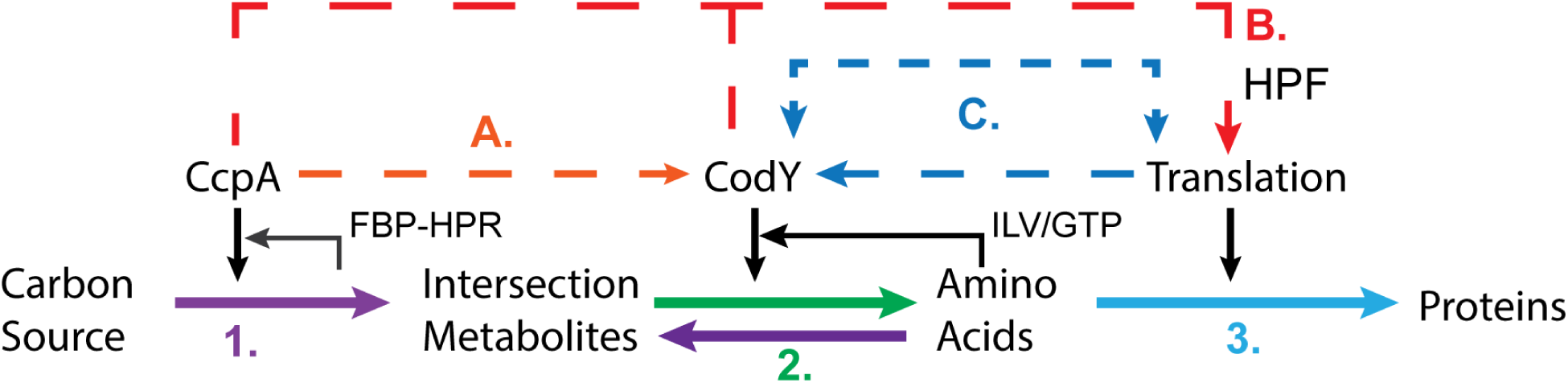
Proposed coarse-grained model of protein biosynthesis regulation in *S. aureus*. The solid lines represent the parts of the protein synthesis pathway controlled by CcpA (purple) and CodY (green). The dashed lines represent new proposed roles of these regulators in (A) coordinating carbon and nitrogen metabolism and (B,C) linking metabolic gene expression with expression of translation associated proteins.

Our analysis suggests that *S. aureus* USA300 strains coordinate their CcpA and CodY activity to regulate carbon and nitrogen flow through the system (dashed orange arrow). Metabolic modeling in CDMG shows increased flux through amino acid biosynthetic reactions when compared to CDM. The results of this TRN-agnostic metabolic model agrees with the increased CodY activity in CDMG and other glucose containing media. Despite close coordination of metabolic flux at different interfaces between CcpA and CodY reactions, it is still not clear how CcpA and CodY activities are coordinated. In *E. coli*, Kochanowski et al. have observed similar coordination between anabolic and catabolic fractions of metabolism^46^. The authors attributed active regulation by Crp and passive changes in metabolic fluxes in response to change in metabolite concentrations as the source of the coordination. Additionally, we also found a feed-forward regulation whereby CcpA and CodY control the expression of the gene encoding HPF protein, which sequesters ribosomes into inactive 100S forms, suggesting a mechanism by which translation is coordinated with metabolic state of the cell (dashed red arrows) ^33,35^.

Lastly, the activity of Translation iModulon is also closely correlated with CodY activity, which may act as an additional layer of coordination between metabolism and translation (dashed blue arrows). However, we have yet to identify the signal or regulator controlling the Translation iModulon activity and therefore the source of this concomitant change in expression along with CodY is unclear. Ribosomal RNA (rRNA) expression is regulated by ppGpp during stringent response which can be activated by mupirocin treatment^28,47^. We therefore expected mupirocin to also have effect on Translation iModulon activity, but we found that while CodY activity increased in response to mupirocin as expected, there was minimal change in Translation activity (**Figure 4c**). This suggests that stringent response, at least when induced by mupirocin treatment, does not play a major role in expression of Translation iModulon genes.

The analysis of the coarse-grained model of metabolic gene regulation presented here was enabled by a computable model of TRN. iModulons enable us to query the TRN at multiple-scales, giving insights into TRN from single gene membership level to global coordination of regulators. By modularizing the TRN, our analysis enabled us to unravel complex regulatory and metabolic interactions to understand regulation of central metabolism one regulator at a time. This modularization can also be used to continually expand on the presented model. For example, our previous works have shown that Translation iModulon activity in *E. coli* and *S. aureus* are closely correlated with stress associated alternate sigma factors^20,21^. This points to a possible entry-point for incorporation of general stress response with metabolism and protein synthesis. Similarly, we have also found that both PyrR and PurR activity is correlated with CodY and CcpA which may provide insights into regulation of nucleotide biosynthesis in response to carbon or nitrogen availability. While we mainly focused on 3 iModulons-CcpA, CodY and Translation-the current model contains 76 total iModulons, each of them rich with information about transcriptional regulation and physiology of *S. aureus*. We thus provide a conceptual framework for overall coordination of metabolism in *S. aureus* and approaches to systematically expand and detail the model proposed.

## Methods

### Strains and Growth Conditions

The *S. aureus* USA300 isolate LAC or its derivative JE2 were used to collect the new RNA sequencing data in this study. The complete description and condition for each of the samples can be found in the model sample table. For RNA sequencing from knock samples, isolated from the Nebraska Transposon Mutant Library were utilized^48^. Unless specified otherwise, samples were grown in duplicates in 20mL of respective media until they reached the O.D_600nm_ of 0.5. 3 mL of culture was harvested and immediately mixed with 6 mL of Qiagen RNA-protect Bacteria Reagent, and incubated at room temperature for 5 minutes. The supernatant was decanted after the samples were centrifuged for 10 mins and 17,500 RPM. The remaining cell pellets were stored in -80C until they were prepared for RNA extraction.

### RNA extraction and sequencing

Total RNA was isolated from the cell pellet in the Qiagen RNeasy Mini Kit columns by following vendor procedures. An on-column DNase treatment was performed for 30 min at room temperature. The ribosomal RNA was removed using RiboRid protocol, as described before^49^. RNA was quantified using a Nanodrop and quality assessed by running an RNA nano chip on a bioanalyser (Agilent, CA). A Swift RNA Library Kit was used following the manufacturer’s protocol to create sequencing libraries.

### Preparing RNA sequencing data for iModulon calculation

The iModulons were calculated from publically available RNA sequencing data from SRA and the newly collected data in this study using pymodulon python package ^41^. The steps used to calculate the iModulons described here were all completed using this package. All RNA sequencing data labeled with *S. aureus* taxonomic ID was downloaded and manually curated to obtain only the samples that were from USA300 isolates. Raw fastq files from curated samples were downloaded, trimmed with TrimGalore and were then aligned to the USA300 TCH1516 genome (NC_010079, NC_012417, NC_010063) using Bowtie2 ^50^. QC/QA stats were collected on each sample using MultiQC and samples that did not pass the QC thresholds (e.g. low read depth, low correlation between replicates, missing metadata) were discarded^51^. Transcripts per million (TPM) was calculated from the remaining high quality RNA sequencing samples. TPM were log transformed and normalized to a control condition within the same BioProject.

### Calculating iModulons from RNA sequencing data

Scipy’s implementation of FastICA was applied to log transformed and normalized TPM data to generate independent components (ICs) and their activities^52,53^. Unlike other decomposition methods, ICA requires the number of dimensions to be calculated as an input. Therefore, various models with different dimensionality were created and the one that maximized regulatory iModulons and minimized single gene iModulon was chosen^54^. The iModulons were then automatically annotated if they overlapped significantly with a curated list of known or predicted regulons and genomic features (e.g. prophages, SCCMec, ACME etc) in *S. aureus*. Other iModulons such as ‘Translation’ or ‘Autolysin’ were manually annotated as all genes contained within the iModulons have a single function.

### Genomic scale modeling of *S. aureus* USA300 metabolism

USA300 specific Genome scale model (GEM) iYS854 was used for all metabolic simulations in the paper. Exchange rate of amino acids, glucose, ammonium and acetate were adjusted to constrain the model to CDM or CDMG specific conditions as described in detail before^29^. Briefly, the uptake or secretion rate for each metabolite from Halsey et al. were normalized by growth-rate, to get growth adjusted solute uptake rate^10^. The exchange rates were then constrained to +/-15% of uptake and exchange rate to account for variance in the data.

Once constrained the model was then used to calculate flux each media using pFBA as implemented in the cobrapy package^30,31^. To get CodY iModulon specific flux, genes in the CodY iModulon were first mapped to metabolic reactions using gene product rule (GPR). The absolute value of fluxes from the pFBA solution for the CodY reactions were then summed to get the final CodY iModulon flux.

To calculate valid amino acid biosynthesis pathway specific flux distribution, the solution spaces of CDM and CDMG specific models were sampled 10,000 times using the Artificial Centering Hit-and-Run algorithm ^55^. Next, the reactions in each amino acid biosynthetic pathway was determined with the MinSpan algorithm^56^. Minspan calculates the set of shortest metabolic pathways that are linearly independent of one another and span the null space of the input model. Each independent pathway defines a mass balanced set of reactions and therefore enables unbiased modularization of metabolism into biologically meaningful pathways. The sampled fluxes (**v**) can therefore be represented as linear weightings (**α**) of minspan pathways (**P**).

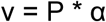

The sampled fluxes were converted to pathway specific weightings (pathway fluxes) using the minspan matrix. Pathways containing amino acid biosynthesis were manually curated and only amino acid biosynthesis pathways that did not appear in multiple MinSpan pathways were used for analysis as they can be easily interpreted and does not require analyzing linear combinations of multiple pathways.

Lastly, the interface metabolites were determined by comparing all metabolites that were involved in at least one CodY and one CcpA reaction. The common metabolites ADP, ATP, CO _2_, coenzyme A, H_2_O, hydrogen atom, sodium ion, NAD, NADH, NADP, NADPH, ammonium (NH4), and phosphate were excluded from this designation.

### Motif enrichment

The 150 base-pairs upstream of *hpf* gene (USA300HOU_RS04065) was scanned for CcpA motif (WTGNNARCGNWWWCAW) using Find Individual Motif Occurence (FIMO) within the MEME suite^39,57^.

## Acknowledgements

This work was funded by the NIAID grant U01AI124316 and the Novo Nordisk Foundation Grant number NNF20CC0035580.

## Data and Code Availability

All RNA-sequencing data used in this work is available publicly on Sequence Reads Archive (SRA). The accession numbers for individual samples can be found in the model sample table. The code used to create the model and generate all the figures in the paper can be found on github (https://github.com/sapoudel/metabs-paper-code).

